# Synchrony of Bird Migration with Avian Influenza Global Spread; Implications for Vulnerable Bird Orders

**DOI:** 10.1101/2023.05.22.541648

**Authors:** Qiqi Yang, Ben Wang, Phillipe Lemey, Lu Dong, Tong Mu, R. Alex Wiebe, Fengyi Guo, Nídia Sequeira Trovão, Sang Woo Park, Nicola Lewis, Joseph Tsui, Sumali Bajaj, Yachang Cheng, Luojun Yang, Yuki Haba, Bingying Li, Guogang Zhang, Oliver G. Pybus, Huaiyu Tian, Bryan Grenfell

## Abstract

Highly pathogenic avian influenza virus (HPAIV) A H5 clade 2.3.4.4 has caused worldwide outbreaks in domestic poultry, occasional spillover to humans, and increasing deaths of diverse species of wild birds since 2014. Wild bird migration is currently acknowledged as an important ecological process contributing to the global dispersal of HPAIV H5. However, it is unclear *how seasonal bird migration facilitates global virus dispersal*, and *which avian species are exposed to HPAI H5 clade 2.3.4.4 viruses and where*. To shed light on ongoing global outbreaks, we sought to explore these questions through phylodynamic analyses based on empirical data of bird movement tracking and virus genome sequences. First, based on viral phylogeography and bird migration networks, we demonstrate that seasonal bird migration can explain salient features of the global dispersal of clade 2.3.4.4. Second, we detect synchrony between the seasonality of bird annual cycle phases and virus lineage movements. We reveal the differing vulnerable bird orders at geographical origins and destinations of HPAIV H5 lineage movements. Notably, we highlight the potential importance of relatively under-discussed Suliformes and Ciconiiformes, in addition to Anseriformes and Charadriiformes, in virus lineage movements. Our study provides a phylodynamic framework that links the bird movement ecology and genomic epidemiology of avian influenza; it highlights the importance of integrating bird behaviour and life history in avian influenza studies.

## 1 Introduction

The re-emergence of highly pathogenic avian influenza viruses (HPAIVs) subtype H5 clade 2.3.4.4 since 2014 [1] has caused unprecedentedly large numbers of wild bird deaths worldwide [2]. In contrast to previous clades of the A/goose/Guangdong/1996 (Gs/GD) lineage, there have also been more persistent spillovers to local domestic poultry [3–7], impacting the poultry farming industry. Despite no onward human-to-human transmission to date, occurrences of zoonotic jumps to humans pose potential threats to public health [8–12]. The unique epidemiological pattern of clade 2.3.4.4 HPAIV H5 is potentially shaped by eco-evolutionary processes: i) the continued selection for both higher transmissibility and virulence, e.g., as observed in ducks [13, 14]; ii) the interaction between the viruses and a wider range of hosts [15].

To shed light on the underlying eco-evolutionary processes, it is critical to understand the spatial dynamics of clade 2.3.4.4 and the ecological factors that influence these patterns. Plausible ecological mechanisms for the global movement of HPAIVs include the live poultry trade and wild bird migration [16, 17]. Preceding the recent re-emergence of clade 2.3.4.4, there has been a long-term debate about whether wild bird migration drives HPAIV dispersal [18, 19]. However, the re-emergence of clade 2.3.4.4 continues to provide virological, epidemiological, and ecological evidence in support of the critical role of migratory wild birds in HPAIV spread and evolution at a global scale. Compared to previous clades, clade 2.3.4.4a during 2014/15 outbreaks was less pathogenic to some species while being more effectively transmitted [20–22], possibly enabling infected birds to migrate between continents. Subsequent phylodynamic work confirmed that the introduction of clade 2.3.4.4a into Europe and North America was most likely via long-distance flights of infected migratory birds [23]. During the 2016/17 outbreaks, the major circulating clade 2.3.4.4b was more transmissible [24] and more virulent [14], related to multiple internal genes [14, 24] and potentially their frequent reassortments [4–6]. Later phylogenetic analysis showed a clear link between the reassortments and migratory birds, as most reassorted gene segments were from migratory wild birds and originated at dates and locations that corresponded to their hosts’ migratory cycles [25]. Integrating host movement in studying HPAIV dispersal is important while challenging. One challenge is insufficient bird movement data, which causes that previous global-scale studies [23] cannot account for the high variation in bird behaviours across species and locations.

Another challenge of studying HPAIV dispersal in wild birds is the lack of HPAIV prevalence data. Only a few studies document longitudinal HPAIV prevalence in wild bird populations[26]. Compared to HPAIV, low pathogenic AIV (LPAIV) has better long-term surveillance of infections or seroprevalence and related avian host ecology in disparate bird habitats, e.g., the United States Geological Survey (USGS) surveillance of birds in Alaska [27]. While longitudinal records provide insights into the role of life history and ecology of local bird communities in LPAIV circulation[28], their conclusions are limited to local dynamics and cannot be easily generalized. To resolve this challenge, ideally, we should have systematic global surveillance for HPAIV. However, this is impossible due to resource constraints.

Instead, we could design effective surveillance strategies by identifying vulnerable avian species and high-risk geographical regions. Recently, researchers have addressed these questions at a higher taxonomic level to include more diverse species. For example, Hill et al. compared the different roles of species within the Anseriformes and Charadriiformes in the dispersal and spillover of AIVs [29]. They concluded that wild geese and swans are the main source species of HPAIV H5, while gulls spread the viruses most rapidly. Hicks et al. found that the inter-species transmission of AIVs in North America is positively associated with the overlap of habitats, suggesting the importance of local bird community diversity [30]. However, they did not use empirical bird movement data. Furthermore, given the heterogeneous biogeographical pattern of bird migrations, identifying geographical hotspots requires linking global and local scales.

To fill this gap, we here focus on two questions related to the contributions of birds, locally and globally, to the spatiotemporal dynamics of HPAIV H5 viruses; specifically, i) how does seasonal bird migration facilitate global virus dispersal and ii) which avian species are exposed to HPAIV H5 and where? To explore these questions, we first illustrate the global circulation history of clade 2.3.4.4 using time-scaled phylogeographic analyses of hemagglutinin (HA) genes of HPAIVs sampled from wild birds and poultry between 2007 and 2018. There are two caveats: first, while we only included HA, internal genes also contribute to virus evolution, e.g., via reassortment [25]; second, the geographical bias of virus sampling has a strong impact on the virus lineage movement routes, especially for locations under-sampled. Based on inferred virus dispersal history, we quantify the contribution of seasonal bird migrations to global virus dispersal and evolution. Second, we model the monthly geographical distribution of bird orders using species distribution models based on environmental factors and bird tracking data. We evaluate the risks of bird orders being exposed to HPAIV H5 at geographical origins and destinations of virus lineage movement by analyzing the statistical association of local bird distributions and virus lineage migration. Our study provides an approach that integrates bird migration ecology in HPAIV epidemiological studies to disentangle the mechanisms of interaction between HPAIV and wild birds.

## 2 Results

### 2.1 Seasonal bird migration associates with global HPAIV H5 dispersal

*Is the wide geographical range of HPAIV H5 clade 2.3.4.4 caused by frequent introductions from one region to another, or a single introduction resulting in subsequent spread within the area?* The discrete-trait phylogeographical analysis of HA genes exhibits scarce virus lineage movements between aggregated regions, most of which are unidirectional (Figure 1A). It suggests that inter-regional viral introductions over long geographical distance occur at low frequency and in one direction. Furthermore, the sequences are highly clustered by region, implying viral persistence within each region after introduction. These patterns qualitatively match bird migration patterns: migratory birds can fly long distances during their migrations, and only fly in one direction in a given season. After arriving at stopover, breeding, or wintering sites, they usually stay for some time, allowing viral transmission to other species or the environment.

**Figure 1:**
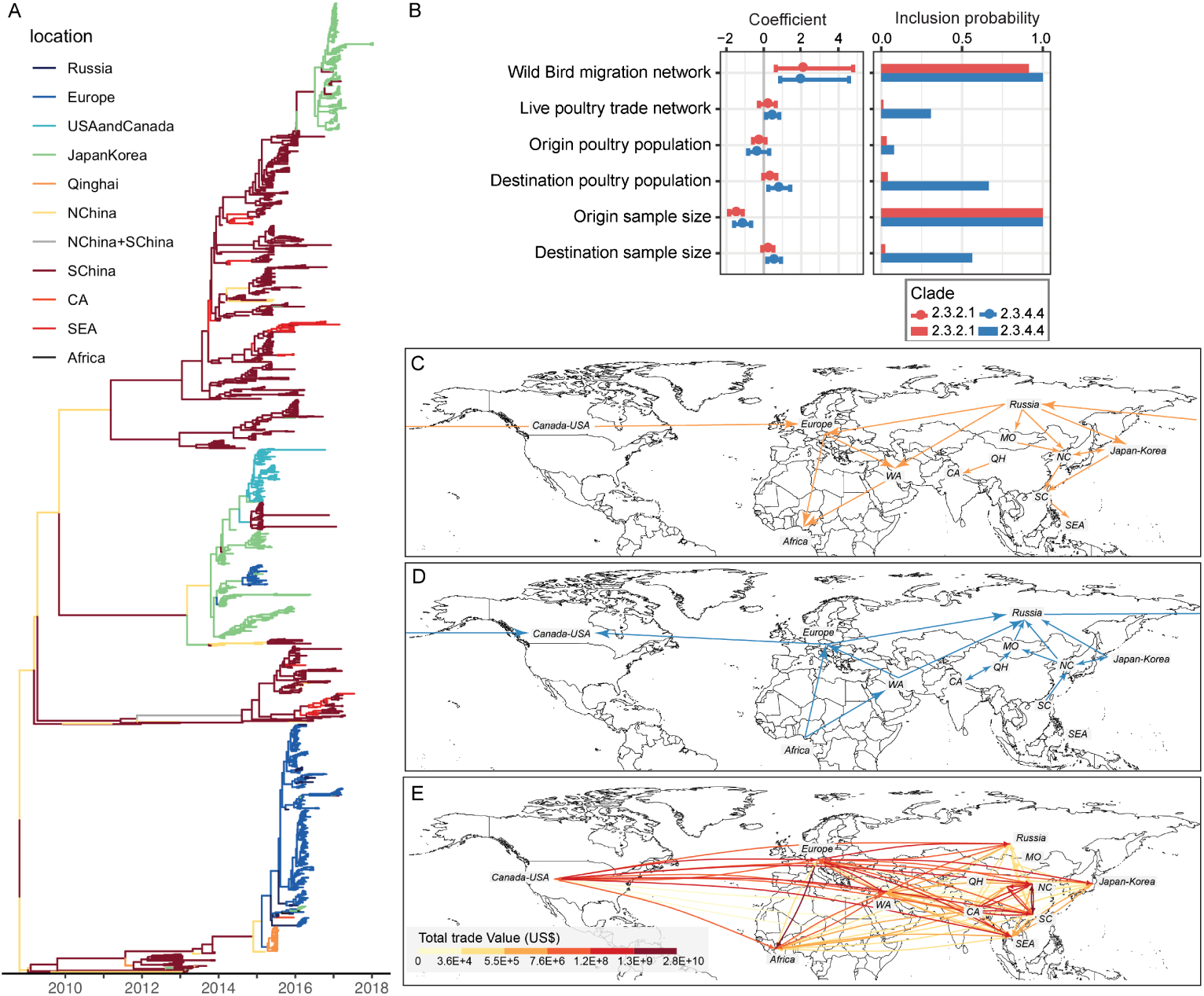
(A) Maximum clade credibility (MCC) time-scaled phylogeny of clade 2.3.4.4 with branches annotated with the inferred location. (B) Contributions of predictors to worldwide diffusion of H5N1 clade 2.3.2.1 and clade 2.3.4.4 inferred from HA genes by GLM-extended Bayesian phylogeographic inference with heterogeneous evolutionary processes through time. Predictors in the model included bird migration network during (C) Northern Hemisphere fall season and (D) Northern Hemisphere spring season, where directed non-weighted edges represent the occurrence of bird migration based on empirical data, and (E) live poultry trade network, where directed weighted edges represent poultry trade value. NChina/NC: North China; SChina/SC: South China; SEA: South-East Asia; CA: Central Asia; QH: Qinghai; MO: Mongolia; WA: Western Asia.

To test quantitatively whether seasonal bird migration is a key predictor of HPAIV H5 dispersal, we fit a generalized linear model (GLM) parameterization of the discrete phylogeography using a Bayesian model selection procedure [31, 32]. Concurrently, we consider seasonal bird migration, live poultry trade and poultry population size as covariates of the diffusion rates between regions. To incorporate the potential seasonal difference in viral dispersal, we model a time-heterogeneous phylogenetic history [33] with three seasons based on bird annual cycle in North Hemisphere: non-migration (mid-November to mid-February, mid-May to mid-September), spring migration (mid-Feburary to mid-May) and fall migration (mid-September to mid-November). Figure 1B shows the posterior estimates of the inclusion probabilities and conditional effect sizes (on a log scale) of the covariates. It reveals that seasonal bird migration is the dominant driver of the global virus lineage movements of HPAIV H5. This is shown in both the log conditional effect size of the seasonal bird migration (mean: *~* 1.96; 95% highest probability density interval, HPDI: 0.88-4.56) and the statistical support for its inclusion (posterior probability *>* 0.999 and Bayes factor *>* 16565).

In contrast, poultry population size and the live poultry trade are not associated with the inter-region dispersal of HPAIV H5 (Figure 1B) in this analysis. It is also evident in both the effect size and the statistical support, e.g, the log conditional effect size of live poultry trade (mean: *~*0.44, HPDI: 0.12-0.84) and the statistical support for its inclusion (posterior probability: *~*0.31 and Bayes factor: 5). To maintain genetic diversity in our data set, we did not down-sample the sequences, which leaves considerable heterogeneity in sample sizes among locations. Therefore, we included the sample size as a predictor in the model to raise the credibility that the inclusion of other predictors is not due to sampling bias. Based on these results, we used subsequent analyses to understand the importance of different bird species at order taxonomy level in the global dispersal and local emergence of HPAIV H5 clade 2.3.4.4.

### 2.2 Vulnerable migratory bird orders at origin and destination regions of HPAIV H5 virus lineage movement

We identified 20 virus dispersal routes (Bayes factor *>*3) between the aggregated regions in the Northern Hemisphere (Figure 2A) using the previous phylogeography analyses. Seasonality is reflected in northward and southward virus lineage movements. Furthermore, it overlaps well with the bird migration seasonality. Most virus lineage movements (14 of 20) show a single temporal peak (Figure S5, 2A). The peaks of the northward routes overlap with spring bird migration and/or wintering period (upper rows of Figure 2A, Figure S5.1). Only one route (Japan-Korea to USA-Canada) overlaps with the summer breeding period. Most southward virus lineage movements peak during the Fall bird migration period, although some peaks continue in November when birds might still be migrating along some routes (lower half of Figure 2A, Figure S5.2). Only one route (Europe to Qinghai) overlaps with the wintering period. In summary, in the Northern Hemisphere, virus lineage movements from south to north occur mainly during the wintering period and spring bird migration, while southward virus lineage movements occur mainly during the fall migration period when birds fly to the south. This association of seasonality in bird migration and HPAIV H5 lineage movement suggests that bird migration is a mechanism of HPAIV H5 global dispersal. It also implies that breeding grounds are potential genetic pools of HPAIV H5 diversity for southward virus lineage movements associated with bird migration; wintering grounds play a similar role for the northward viral lineage movements. Additionally, the results show more virus lineage movements during the fall migration (Southward Markov Jump counts: 310498 per month, September-November) than the spring migration (Northward Markov Jump counts: 257503 per month, March-May). Virus lineage movements also have higher relative frequency during the fall migration (shown in the higher peak in Figure 2). Interestingly, birds also migrate in a larger abundance in the fall than during spring, as the population size becomes larger after breeding.

**Figure 2:**
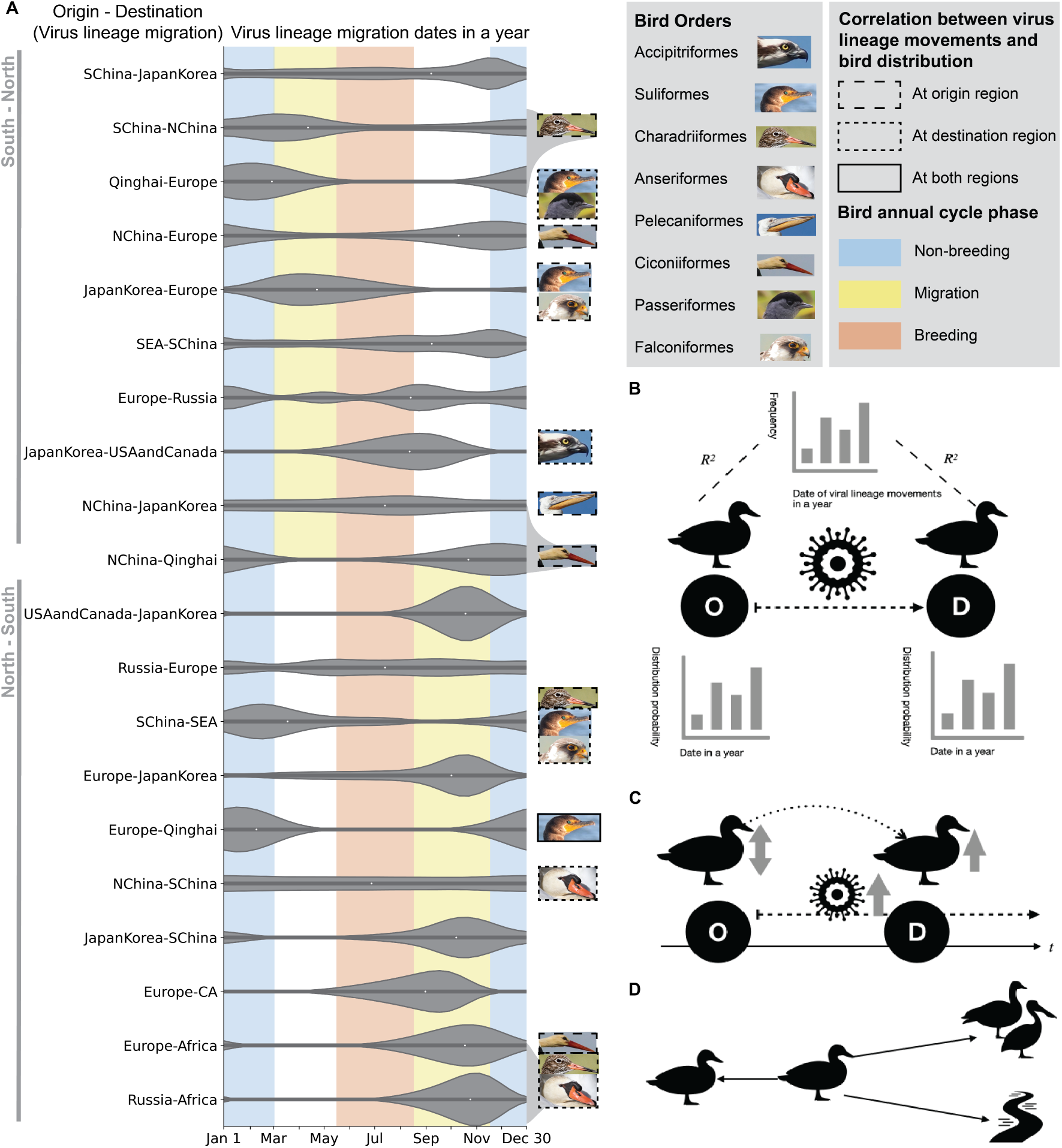
(A) Probability density distribution of the virus lineage migration throughout the year, between locations summarized from the discrete trait phylogeography of HPAIV H5 clade 2.3.4.4 and the Markov jump counts (Section 4.3). *X* axis: Virus lineage migration dates in a year; labels on *Y* axis: origin region - destination region of the virus lineage migration. The width of the violins represents the virus lineage migration probability density. Boxes around bird photos show the statistically significant correlation of virus lineage movements and bird order distribution at origin, destination or both regions. Bird species photos were obtained from the Macaulay Library at the Cornell Lab of Ornithology (macaulaylibrary.org). The entries of the photos are listed in Table S5. Non-breeding (blue), migration (yellow) and breeding (red) bird annual cycle phases in general are shown in the south-north migration direction and in the north-south migration direction. (B) Schematic diagram of cross-correlation analyses of virus lineage movement between two locations (O: origin, D: destination) and the bird distribution probability at each location. (C) Time scale of virus lineage movement, bird migration and local virus transmission, including inter-species, inter-individual and environmental transmissions. The grey arrows indicate the increase or decrease of the local bird population and the virus lineage movement influx. (D) Local transmission of AIV includes inter-individual transmission within a population, inter-species transmission within a bird community and potential environmental transmission.

*Which migratory bird orders might be exposed to HPAIV H5 at the origin or destination regions of virus lineage movements?* To explore this question, we examined the synchrony of bird order distribution and virus lineage movements. The result shows that 8 bird orders (out of 9) at origin or destination regions are correlated with 12 virus lineage movement routes (out of 20) (Table S4, Figure 2A). Notably, the distribution of Suliformes, e.g., cormorants, during a year in Europe (*r* = 0.996, 95% confidence intervals, CI: [-0.566, 0.566], *p <* 0.001) and Qinghai (*r* = 0.899, CI: [-0.566, 0.566], *p <* 0.002) synchronizes with virus lineage movements from Europe to Qinghai, suggesting that Suliformes might be associated with HPAIV H5 spread from Europe to Qinghai. However, due to the possible under-sampling of viruses in northern and central Eurasia, we cannot conclude that virus lineage movements occur directly from Europe to Qinghai. In addition, Suliformes, along with Charadriiformes, Ciconiiformes, and Anseriformes, are associated with multiple (*>*2) virus lineage movements. Three routes of virus lineage movement are related to the distribution of multiple (*>*2) bird orders:

- The virus lineage movement from Qinghai to Europe is associated with the Charadriiformes distribution in Qinghai (*r* = *−*0.820*, p <* 0.005), and the distribution of Suliformes (*r* = 0.924*, p <* 0.001) and Passeriformes (*r* = 0.878*, p <* 0.002) in Europe.
- The virus lineage movement from South China to South East Asia synchronizes with the Charadriiformes distribution in South China (*r* = 0.803, *p <* 0.005), and the distribution of Suliformes (*r* = 0.912*, p <* 0.002) and Falconiformes (*r* = 0.890*, p <* 0.002) in South East Asia.
- The virus lineage movement from Europe to Africa is related to the Ciconiiformes distribution in Europe (*r* = 0.813, *p <* 0.005), and the distribution of Charadriiformes (*r* = 0.886, *p <* 0.002) and Anseriformes in Africa (*r* = 0.905, *p <* 0.002).

Despite the possible geographical sampling bias, our results suggest integrating host distribution inference and phylogeographic analysis might be able to retrospectively identify important bird species and geographical regions in avian influenza transmission.

## 3 Discussion

Here, we report a phylodynamic analysis linking spatial ecology of avian hosts and HPAIV H5 virus lineage movements. Our results support previous findings on the important role of bird migration in the dissemination of HPAIV H5 clade 2.3.4.4 [23]. We found that the seasonal wild bird migration network is associated with the global diffusion and evolutionary dynamics of HPAIV H5. A previous study found that the 2014/2015 outbreaks of HPAIV H5 in Europe and North America were likely introduced by wild bird migration [23] by comparing the inferred ancestral host-type and location traits of the viral genome sequences [23]. Our study advances this finding by directly integrating the bird migration trajectory network into the virus phylogeographic reconstruction. In addition, we found that inter-regional live poultry trade is not associated with the global HPAIV H5 dispersal, consistent with previous studies [16, 23]. The same previous study found that the international poultry trade’s direction is opposite to the global spread direction of HPAIV H5 clade 2.3.4.4 [23]. Another previous study demonstrated largescale H5N1 transmission dynamics are structured according to different bird flyways and driven by the Anatidae family, while the Phasianidae family, largely representing poultry, is an evolutionary sink [16].

Historically, Anseriformes have been the focus of wild bird hosts when studying host-pathogen interaction in AIV studies. However, many other understudied orders have been affected by clade 2.3.4.4 recently [15, 34]. Interaction of different avian orders might contribute to virus dispersal and local persistence [29]. A previous study showed that host origins of HPAIV H5 reassorted genes include Anseriformes, other groups of wild birds, and domestic poultry [25].

*Caveats*. A limitation of our results is that the undersampling of viruses in some areas hugely impacted the inferred phylogeography. For example, we cannot conclude if the inferred viral lineage movement from Europe to Qinghai or Japan-Korea occurs directly or if geographically-proximate areas, e.g., central Eurasia, are middle stops of the movement, due to under-sampling in central Eurasia. Despite including sampling size in the phylogeographical analysis, we cannot adjust the geographical sampling biases due to the unknown magnitude of infections at locations. Fortunately, the sampling efforts in some historically under-sampled and no-sampled areas are growing, e.g., in Australia [35]. In the future, given more extensive and evenly-sampled spatial data, our methods could be utilized to understand the role of wild birds in virus dispersal.

Despite using empirical bird movement data, our analyses include limited species diversity and dispersal area. Therefore, we did not include the migration volume of birds in the migration network (Figure 1). Currently, bird migration is summarized as a binary network. In the future, integrating comprehensive bird movement models [36, 37] would provide a more detailed understanding of the mechanism of how bird migration contributes to AIV dispersal. Another caveat is that we only considered the HA gene when inferring AIV diffusion and evolution. HA is a key gene in influenza viruses, as it is the receptor-binding and membrane fusion glycoprotein of influenza virus and the target for infectivity-neutralizing antibodies [38]. However, the reassortment events of all internal genes are also important in the dispersal and evolution of HPAIV H5 [25].

Our results also show high spatial and temporal heterogeneity in the association strength between specific bird orders and virus lineage movements. Despite the low relative frequency of virus lineage movements during summer breeding and wintering, they may serve as a gene pool for following virus lineage movement during the migration. A previous study emphasizes the important role of the breeding period in interspecies virus transmission in North America [30]. Previous surveillance also shows that LPAI prevalence in waterfowls is higher during the wintering period of Eurasian migratory birds in Africa [39]. Additionally, our results highlight the importance of Suliformes and Ciconiiformes in HPAIV H5 dispersal, which are understudied compared to Anseriformes and Charadriiformes.

We did not account for possible interspecies transmission among individuals of multiple bird orders. This is a possible reason for associations between some bird orders and virus dispersal routes where there is no direct bird migration between the origin and destination location. For example, the spring migration of Suliformes and Falconiformes overlaps with virus lineage movements from Japan-Korea to Europe. While birds might not directly fly between the two regions, various species stop between Japan and Europe during migration. Interspecies transmission at the stop-over sites might lead to the virus lineage movements (Figure 2D). However, the under-sampling of the viruses and lack of bird tracking data might also contribute to the observed pattern.

Another limitation is that we did not account for variation in movement behaviour within each bird order. Due to limited data, bird order is the most accurate taxonomy level we can study reliably. Finally, we included virus samples from domestic poultry when inferring virus diffusion. Therefore, some patterns in the results could reflect virus transmission between domestic poultry and spillover from wild birds to poultry rather than bird migratory patterns.

In conclusion, allocating more resources for global surveillance of avian influenza viruses in wild birds would enhance our ability to tackle the challenges of more virulent and transmissible HPAIV H5 spreading in wild birds. To achieve this goal, it is critical to understand “where and in which bird species surveillance is most needed and could have the greatest impact” [17]. Given sufficient data in the future, our framework could help conservation and public health policy-making in designing monitoring and surveillance strategies. More collaboration is needed between ornithologists, movement ecologists, bird conservation experts, avian influenza epidemiologists, disease ecologists and virologists on many aspects, including collaborative data collection/surveillance of AIV and data sharing. For example, if studies were to simultaneously obtain the movement tracking of bird populations and their serology and virology surveillance data, then they could link the bird movement directly with the virus transmission and dispersal. In addition, we need more AIV samples from water bodies to better understand environmental transmission. With such data, we would be able to understand the viral transmission at local scales and therefore develop disease models for bird conservation and potential zoonotic threats.

## 4 Materials and Methods

### 4.1 Wild bird movement tracking and distribution modeling

To assemble the global wild bird observation data, we accessed the worldwide bird tracking data from Movebank in 2021. This dataset amassed from 53 studies across the world [40–119]. The Movebank study ID, name, principal investigator, and contact person are listed in Table S6. The dataset is collected by various research groups, and by various sensors, including Global Position System (GPS), Argos, bird ring, radio transmitter, solar geo-locator, and natural mark. It covers over 3542 individual birds (class: Ave), including 10 orders and 95 species (Table S1). For further modelling the migration of the wild birds belonging to different orders, we excluded the observation data on Movebank of Cuculiformes, Caprimulgiformes, Strigiformes, Columbiformes, Phoenicopteriformes, Piciformes, Sphenisciformes, and Procellariiformes, given their paucity and geographically restricted distribution. Additionally, we accessed GPS tracking data of 193 individuals, including 5 orders and 12 Species between 2006 and 2019 in China from a previous study ([120]). Accordingly, we combined the data from China with those on Movebank (Table S6) and finalized a bird observation dataset consisting of 10 orders and 96 species.

To model the wild bird distribution throughout a year, we developed a model framework based on the species distribution model (SDM). The response variable of the model is bird occurrence (1: presence; 0: pseudo-absence). The independent variables are 20 well-studied environmental predictors, including local topography, weather conditions, and time of the season. Table S2 lists the environmental data and the source. We divide the globe into 1-km resolution geographical cells for each month. For each cell, the value of the dependent variable is 1 if there is any observation of an individual in the target order in that month in the bird tracking data, otherwise 0. Furthermore, to infer the probability of bird occurrence between 0 and 1 for each cell, we trained a XGBoost binary classification model [121] for each bird order, respectively. The method is adapted from a previous bird migration model [122]. We used true presence and pseudo-absence data (marked as 1 and 0 respectively). We fitted the distribution of birds which manifest as true-presence data and pseudo-absence data. We randomly divided 67% of the data as the training set and the other 33% as the test set. The model finally outputs the probability of the distribution of migratory birds in each month across years (Dataset 6). The accuracy was evaluated by the area under the curve (AUC) in a test set of the ten orders: Pelecaniformes (0.97), Gruiformes (0.97), Passeriformes (0.97), Suliformes (0.98), Ciconiiformes (0.92), Falconiformes (0.98), Charadriiformes (0.94), Anseriformes (0.90), Accipitriformes (0.90). The modelled wild bird distribution (Dataset 6) was applied in the subsequent analysis to identify key bird orders associated with the global viral dispersal (Section 4.3) and local virus emergence.

### 4.2 Viral sequence data and time-scaled phylogeny of HPAIVs

To infer the phylogeny of avian influenza HPAIV H5 viruses, we accessed sequences of HA genes, NA genes and six internal gene segments from GISAID (Global Initiative on Sharing All Influenza Data [123–125]). Using the sequences, we estimated a maximum likelihood phylogeny (Figure S3) for each gene segment, respectively, under a GTR+*γ* nucleotide substitution model, with the randomly selected strains as representatives, by FastTree v2.1.4 [126]. Genotypes of internal gene segments (Figure S3) were defined by clustering pattern with background sequences in a previous study [127]. On the phylogeny, the viruses with internal genes from wild birds, e.g. clade 2.3.2.1 and clade 2.3.4.4, showed wider geographical spread [1, 23], compared to poultry viruses, e.g. clade 2.3.4.1 and clade 2.2, despite the high similarity of their HA genes. This demonstrates the importance of gene reassortment in the evolution and transmission of HPAIVs. In this project, we focused on clade 2.3.4.4 and clade 2.3.2.1. Next, we inferred their time-scaled phylogenies of HA genes. Before the inference, to test for the presence of phylogenetic temporal structure, we generated a scatter-plot of root-to-tip genetic divergence against sampling date by TempEst v1.5 [128]. Strong phylogenetic temporal structure was detected in the phylogeny of each clade (Figure S7). The final datasets (Dataset 2) were i) 1163 HA sequences of clade 2.3.2.1 ii) 1844 HA sequences of clade 2.3.4.4. The spatial and temporal distribution of the sequences is shown in Figure S4.

Time-resolved HA phylogenies were estimated using the Markov chain Monte Carlo (MCMC) approach implemented in BEAST v1.10.4 [129] with the BEAGLE library [130]. We used an uncorrelated lognormal (UCLN) relaxed molecular clock model [131], the SRD06 nucleotide substitution model [132] and the Gaussian Markov random field (GMRF) Bayesian Skyride coalescent tree prior [133]. For each dataset, MCMC chains were run for 300 million (clade 2.3.2.1) and 400 million (clade 2.3.4.4) generations with burn-in of 10%, sampling every 10,000 steps. Convergence of MCMC chains was checked with Tracer v1.7 [134]. A set of 1000 trees for each clade was subsampled from the MCMC chain and used as an empirical tree distribution for the subsequent analysis.

### 4.3 Discrete trait phylogeography of HPAIVs and counts of virus lineage migration

Based on empirical phylogenies, we used a non-reversible discrete-state continuous time Markov chain (CTMC) model and a Bayesian stochastic search variable selection (BSSVS) approach [31] to infer the viral diffusion among locations: i) the most probable locations of the ancestral nodes in the phylogeny and ii) the history and rates of lineage movement among locations [31]. Sampled countries were divided into 10 locations: Africa, Central Asia, Europe, Japan-Korea, North China, South China, Qinghai, Russia, Southeast Asia and USA-Canada. This regional categorization was done according to the major wild bird breeding areas. Furthermore, to estimate the viral gene flows between locations, we used a robust counting approach [135, 136] to count virus lineage migration events. The basic idea is to count the expected number of lineage movements (Markove jumps) between the locations along the phylogeny branches, as applied in previous studies [137–141]. For each location, the frequency distribution throughout a year of the Markov jumps from or to the place is summarized. Using this method, we summarized monthly frequency distribution of the virus lineage migration for each pathway (Figure S5, Dataset 4). This was used for further analysis below.

To target the key bird orders for each location, we explored the association of wild bird distribution across a year and the virus diffusion. The monthly wild bird distribution probability at each location (Dataset 5) is generated based on the location’s geographical coordinates on the modelled bird distribution probability raster map (Dataset 6). We calculated the correlation between the virus lineage migration and the bird probability distribution at origin and destination regions, respectively, with time lags from −7 to 7. To account for multiple comparisons of 9 bird orders, we use *p* value *<* 0.00556(= 0.05*/*9) to define the statistical significance in the correlations. When bird distribution at the origin leads to the virus lineage movements positively or negatively, we consider the bird order distribution at the origin to be correlated with the virus lineage movements (Table S4.1, Figure 2A). When bird distribution at the sink is positively associated with the virus lineage movement, we consider the bird order distribution at the sink is correlated with the virus lineage movement (Table S4.2, Figure 2A).

### 4.4 Animal mobility networks and their contribution to HPAIV phylogeography

The bird migration network (Figure 1C, D) was summarised by searching publicly available migration data on Movebank. An edge between two locations in the network exists if any migration tracking record shows bird migration. The location-wise live poultry trade values (Dataset 1) were summed up from country-wise import and export of the live poultry recorded on United Nations Comtrade Database (comtrade.un.org/data/). We accessed the total net weight and trade value from 1996 to 2016 of live poultry, including fowls of the species Gallus domesticus, ducks, geese, turkeys and guinea fowls. Since there are no accessible data of the within-country poultry trade in China, we adapted the inferred poultry trade accessibility between provinces of China from a previous study [142]. Based on the ratio of the inferred accessibility and the empirical trade value between Hong Kong SAR and the mainland of China, we scaled all the accessibility to the trade value flows among Qinghai, North China and South China.

With the summarized seasonal-varying bird migration network, we statistically quantified the contribution of wild bird migration to avian influenza virus dispersal. We applied the generalized linear model (GLM) extended Bayesian phylogeography inference [32] with the 1000 empirical trees as the input. The 11 categorized locations in the previous discrete trait phylogeography were still used. The epoch model [33] was used to model the time heterogeneity of the contribution. To explain the contribution of the bird migration and the respective seasonal migration, we also separated the network of spring migration and that of the fall as two predictors for comparison (Figure S6). For each clade and each predictor group, MCMC chains were run for 100 million generations with burn-in of 10%, sampling every 10,000 steps. Similarly, we assessed the convergence of the chains in Tracer v1.7 [134].

## Acknowledgements

We gratefully acknowledge all bird tracking data contributors, i.e., the authors and other researchers in their originating groups collecting the data and metadata and sharing via Movebank, on which this research is based. The principle investigator, contact person, citation or data repository DOI are listed in Supplementary Table S6. We also gratefully acknowledge all data contributors of virus genome sequences, i.e., the authors and their originating laboratories responsible for obtaining the specimens, and their submitting laboratories for generating the genetic sequence and metadata and sharing via the GISAID Initiative, on which this research is based.

We thank Qizhong Wu and Joint Center for Earth System Modeling and High Performance Computing, Beijing Normal University, for providing computing resources for time-scaled phylogenetic analyses. We thank Jing Yu for assisting in assembling international poultry trade data. We thank Bram Vrancken for assisting with phylogenetic analyses. We thank insightful comments from Andrew Rambaut and the Virus Club Group at the University of Edinburgh, Grenfell lab at Princeton University, the Influenza group at Francis Crick Institute, Olga Alexandrou at Society for the Protection of Prespa, Yonghong Liu at Chinese Center for Disease Control and Prevention, Yidan Li (formerly at Beijing Normal University), and Nils Stenseth at the University of Oslo. Y.H. is supported by a Masason Foundation Fellowship, a Honjo International Fellowship, and a Centennial Fellowship. The opinions expressed in this article are those of the authors and do not reflect the view of the National Institutes of Health, the Department of Health and Human Services, or the United States government.

## 5 Data Availability Statement

We provide Movebank Study ID (unique searchable identifier) and relevant metadata information for Movebank bird tracking data. We also provide accession ID for GI-SAID virus genomic data. All code scripts for analyzing data are provided. All data and scripts are available as a public project https://doi.org/10.17605/OSF.IO/7A2UK on Open Science Framework and GitHub Repository https://github.com/kikiyang/HPAI_Bird_world.

## References

1. Smith, G. J. & Donis, R. O. Nomenclature updates resulting from the evolution of avian influenza A(H5) virus clades 2.1.3.2a, 2.2.1, and 2.3.4 during 2013-2014. Influenza and other Respiratory Viruses 9. ISBN: 1750-2640, 271–276. issn: 17502659 (2015).

2. Wille, M. & Barr, I. G. Resurgence of avian influenza virus. Science 376. Publisher: American Association for the Advancement of Science, 459–460. https://www.science.org/doi/10.1126/science.abo1232 (2022) (Apr. 29, 2022).

3. 2020: OIE - World Organisation for Animal Health en. https://www.oie.int/animal-health-in-the-world/update-on-avian-influenza/2020/ (2021).

4. Beerens, N. et al. Multiple Reassorted Viruses as Cause of Highly Pathogenic Avian Influenza A(H5N8) Virus Epidemic, the Netherlands, 2016. Emerging Infectious Diseases 23, 1974–1981. issn: 1080-6040. https://www.ncbi.nlm.nih.gov/pmc/articles/PMC5708218/ (2021) (Dec. 2017).

5. Kwon, J.-H. et al. New Reassortant Clade 2.3.4.4b Avian Influenza A(H5N6) Virus in Wild Birds, South Korea, 2017–18. Emerging Infectious Diseases 24, 1953–1955. issn: 1080-6040. https://www.ncbi.nlm.nih.gov/pmc/articles/PMC6154165/ (2021) (Oct. 2018).

6. Mine, J. et al. Genetics and pathogenicity of H5N6 highly pathogenic avian influenza viruses isolated from wild birds and a chicken in Japan during winter 2017-2018. eng. Virology 533, 1–11. issn: 1096-0341 (July 2019).

7. Lee, D.-h., Bertran, K., Kwon, J.-h. & Swayne, D. E. Evolution, global spread, and pathogenicity of highly pathogenic avian influenza H5Nx clade 2.3.4.4. J Vet Sci 18. ISBN: 1706546343, 269–280 (2017).

8. Update on Avian Influenza A (H5N1) Virus Infection in Humans. New England Journal of Medicine 358. Publisher: Massachusetts Medical Society eprint: https://doi.org/10.10261#273. issn: 0028-4793. https://doi.org/10.1056/NEJMra0707279 (2021) (Jan. 2008).

9. Gao, R. et al. Human Infection with a Novel Avian-Origin Influenza A (H7N9) Virus. New England Journal of Medicine 368. Publisher: Massachusetts Medical Society eprint: https://doi.org/10.1056/NEJMoa1304459, 1888–1897. issn: 0028-4793. https://doi.org/10.1056/NEJMoa1304459 (2021) (May 2013).

10. Yang, Z.-F., Mok, C. K., Peiris, J. S. & Zhong, N.-S. Human Infection with a Novel Avian Influenza A(H5N6) Virus. New England Journal of Medicine 373. Publisher: Massachusetts Medical Society eprint: https://doi.org/10.1056/NEJMc1502983, 487–489. issn: 0028-4793. https://doi.org/10.1056/NEJMc1502983 (2021) (July 2015).

11. First identification of human cases of avian influenza A (H5N8) infection. en, 9.

12. Peiris, J. S. M., de Jong, M. D. & Guan, Y. Avian Influenza Virus (H5N1): a Threat to Human Health. Clinical Microbiology Reviews 20. Publisher: American Society for Microbiology, 243–267. https://journals.asm.org/doi/full/10.1128/CMR.00037-06 (2022) (Apr. 2007).

13. Grund, C., et al. A novel European H5N8 influenza A virus has increased virulence in ducks but low zoonotic potential. Emerging Microbes & Infections 7. Publisher: Taylor & Francis eprint: https://doi.org/10.1038/s41426-018-0130-1, 1–14. issn: null. https://doi.org/10.1038/s41426-018-0130-1 (2022) (Dec. 1, 2018).

14. Leyson, C. M., Youk, S., Ferreira, H. L., Suarez, D. L. & Pantin-Jackwood, M. Multiple Gene Segments Are Associated with Enhanced Virulence of Clade 2.3.4.4 H5N8 Highly Pathogenic Avian Influenza Virus in Mallards. Journal of Virology 95. Publisher: American Society for Microbiology, e00955–21. https://journals.asm.org/doi/10.1128/JVI.00955-21 (2022) (2021).

15. European Food Safety Authority, European Centre for Disease Prevention, Control, European Union Reference Laboratory for Avian Influenza et al. Avian Influenza Overview December 2021 – March 2022. EFSA Journal 20. issn: 18314732, 18314732. https://data.europa.eu/doi/10.2903/j.efsa.2022.7289 (2022) (Apr. 2022).

16. Trovão, N. S., Suchard, M. A., Baele, G., Gilbert, M. & Lemey, P. Bayesian Inference Reveals Host-Specific Contributions to the Epidemic Expansion of Influenza A H5N1. Molecular Biology and Evolution 32, 3264–3275. issn: 0737-4038. https://doi.org/10.1093/molbev/msv185 (2021) (Dec. 2015).

17. Russell, C. A. Sick birds don’t fly…or do they? Science 354. ISBN: 1095-9203 (Electronic) 0036-8075 (Linking), 174–175. issn: 10959203 (2016).

18. Krauss, S. et al. The enigma of the apparent disappearance of Eurasian highly pathogenic H5 clade 2.3.4.4 influenza A viruses in North American waterfowl. en. Proceedings of the National Academy of Sciences 113. Publisher: National Academy of Sciences Section: Biological Sciences, 9033–9038. issn: 0027-8424, 1091-6490. https://www.pnas.org/content/113/32/9033 (2021) (Aug. 2016).

19. Dennis, N. Are Wild Birds to Blame ? Science 310, 426–428 (2005).

20. Kim, Y.-I., et al. Pathobiological features of a novel, highly pathogenic avian influenza A(H5N8) virus. Emerging Microbes & Infections 3. Publisher: Taylor & Francis, 1–13. issn: 2222-1751. https://www.tandfonline.com/doi/full/10.1038/emi.2014.75 (Jan. 2014).

21. Kang, H.-M. et al. Novel Reassortant Influenza A(H5N8) Viruses among Inoculated Domestic and Wild Ducks, South Korea, 2014. Emerging Infectious Diseases 21, 298–304. issn: 1080-6040. https://www.ncbi.nlm.nih.gov/pmc/articles/PMC4313655/ (2021) (Feb. 2015).

22. Sun, H., et al. Characterization of clade 2.3.4.4 highly pathogenic H5 avian influenza viruses in ducks and chickens. Veterinary Microbiology 182. Publisher: Elsevier B.V., 116–122. issn: 18732542. http://dx.doi.org/10.1016/j.vetmic.2015.11.001 (2016).

23. Lycett, S. J. et al. Role for migratory wild birds in the global spread of avian influenza H5N8. Science 354. ISBN: 1095-9203 (Electronic)$\backslash$r0036-8075 (Linking), 213–217. issn: 0036-8075. http://www.sciencemag.org/cgi/doi/10.1126/science.aaf8852 (Oct. 2016).

24. Blaurock, C. et al. Preferential Selection and Contribution of Non-Structural Protein 1 (NS1) to the Efficient Transmission of Panzootic Avian Influenza H5N8 Virus Clades 2.3.4.4A and B in Chickens and Ducks. Journal of Virology 95. Publisher: American Society for Microbiology, e00445–21. https://journals.asm.org/doi/10.1128/JVI.00445-21 (2022) (2021).

25. Lycett, S. J. et al. Genesis and spread of multiple reassortants during the 2016/2017 H5 avian influenza epidemic in Eurasia. Proceedings of the National Academy of Sciences of the United States of America 117, 20814–20825. issn: 10916490 (2020).

26. Hill, S. C. et al. Comparative Micro-Epidemiology of Pathogenic Avian Influenza Virus Outbreaks in a Wild Bird Population. Philosophical Transactions of the Royal Society B: Biological Sciences 374, 20180259. https://royalsocietypublishing.org/doi/10.1098/rstb.2018.0259 (2022) (June 24, 2019).

27. Reeves, A. B., et al. Influenza A virus recovery, diversity, and intercontinental exchange: A multi-year assessment of wild bird sampling at Izembek National Wildlife Refuge, Alaska. PLOS ONE 13. Publisher: Public Library of Science, e0195327. issn: 1932-6203. https://journals.plos.org/plosone/article?id=10.1371/journal.pone.0195327 (2022) (Apr. 5, 2018).

28. Van Dijk, J. G. B. et al. Juveniles and migrants as drivers for seasonal epizootics of avian influenza virus. Journal of Animal Ecology 83. eprint: https://onlinelibrary.wiley.com/doi/2656.12131, 266–275. issn: 1365-2656. https://onlinelibrary.wiley.com/doi/abs/10.1111/1365-2656.12131 (2022) (2014).

29. Hill, N. J., et al. Ecological divergence of wild birds drives avian influenza spillover and global spread. PLOS Pathogens 18. Publisher: Public Library of Science, e1010062. issn: 1553-7374. https://journals.plos.org/plospathogens/article?id=10.1371/journal.ppat.1010062 (2022) (May 19, 2022).

30. Hicks, J. T. et al. Host diversity and behavior determine patterns of interspecies transmission and geographic diffusion of avian influenza A subtypes among North American wild reservoir species. PLOS Pathogens 18. Publisher: Public Library of Science, e1009973. issn: 1553-7374. https://journals.plos.org/plospathogens/article?id=10.1371/journal.ppat.1009973 (2022) (Apr. 13, 2022).

31. Lemey, P., Rambaut, A., Drummond, A. J. & Suchard, M. A. Bayesian Phylogeography Finds Its Roots. PLoS Computational Biology 5 (ed Fraser, C.) ISBN: 1553-734X, e1000520. issn: 1553-7358. https://dx.plos.org/10.1371/journal.pcbi.1000520 (Sept. 2009).

32. Lemey, P. et al. Unifying Viral Genetics and Human Transportation Data to Predict the Global Transmission Dynamics of Human Influenza H3N2. PLoS pathogens 10. ISBN: 1553-7374 (Electronic)$\backslash$r1553-7366 (Linking), 1–17. issn: 1553-7374. http://www.scopus.com/inward/record.url?eid=2-s2.0-84895733259&partnerID=tZOtx3y1 (2014).

33. Bielejec, F., Lemey, P., Baele, G., Rambaut, A. & Suchard, M. Inferring heterogeneous evolutionary processes through time: from sequence substitution to phylogeography. en. Syst Biol 63, 493–504 (2014).

34. Ramey, A. M. et al. Highly pathogenic avian influenza is an emerging disease threat to wild birds in North America. The Journal of Wildlife Management 86. eprint: https://onlinelibrary.wiley.com/doi/pdf/10.1002/jwmg.22171, e22171. issn: 1937-2817. https://onlinelibrary.wiley.com/doi/abs/10.1002/jwmg.22171 (2022) (2022).

35. Wille, M. et al. Australia as a Global Sink for the Genetic Diversity of Avian Influenza A Virus. PLOS Pathogens 18, e1010150. issn: 1553-7374. https://journals.plos.org/plospathogens/article?id=10.1371/journal.ppat.1010150 (2023) (May 10, 2022).

36. Van Toor, M. L., Avril, A., Wu, G., Holan, S. H. & Waldenström, J. As the Duck Flies—Estimating the Dispersal of Low-Pathogenic Avian Influenza Viruses by Migrating Mallards. Frontiers in Ecology and Evolution 6. issn: 2296-701X. https://www.frontiersin.org/article/10.3389/fevo.2018.00208 (2022) (2018).

37. Van Toor, M. L. et al. Integrating animal movement with habitat suitability for estimating dynamic migratory connectivity. Landscape Ecology 33, 879–893. issn: 1572-9761. https://doi.org/10.1007/s10980-018-0637-9 (2022) (June 1, 2018).

38. Skehel, J. J. & Wiley, D. C. Receptor Binding and Membrane Fusion in Virus Entry: The Influenza Hemagglutinin. Annual Review of Biochemistry 69, 531–69. issn: 00664154. https://www.proquest.com/docview/223690008/abstract/D796D35D31C8400FPQ/1 (2023) (2000).

39. Gaidet, N., et al. Understanding the ecological drivers of avian influenza virus infection in wildfowl: a continental-scale study across Africa. Proceedings of the Royal Society B: Biological Sciences 279. Publisher: Royal Society, 1131–1141. https://royalsocietypublishing.org/doi/full/10.1098/rspb.2011.1417 (2022) (Mar. 22, 2012).

40. Warwick-Evans, V., et al. Data from: Changes in Behaviour Drive Inter-annual Variability in the At-sea Distribution of Northern Gannets 2017.

41. Friedemann, G., Leshem, Y. & Izhaki, I. Data from: Multidimensional Differentiation in Foraging Resource Use during Breeding of Two Sympatric Raptors: Space, Habitat Type, Time and Food 2016.

42. Hallworth, M. & Marra, P. Data from: Miniaturized GPS Tags Identify Non-Breeding Territories of a Small Breeding Migratory Songbird 2015.

43. Petersen, M. & Douglas, D. Data from: At-sea Distribution of Spectacled Eiders: A 120-Year-Old Mystery Resolved 2016.

44. Spiegel, O., Getz, W. & Nathan, R. Data from: Factors Influencing Foraging Search Efficiency: Why Do Scarce Lappet-Faced Vultures Outperform Ubiquitous White-Backed Vultures? (V2) 2014.

45. Arlt, D., Olsson, P., Fox, J., Low, M. & Pärt, T. Data from: Prolonged Stopover Duration Characterises Migration Strategy and Constraints of a Long-Distant Migrant Songbird 2015.

46. Kleyheeg, E., van Dijk, J., Nolet, B. & Soons, M. Data from: Movement Patterns of a Keystone Waterbird Species Are Highly Predictable from Landscape Configuration 2017.

47. Exo, K. Data from: Forecasting Spring from Afar? Timing of Migration and Predictability of Phenology along Different Migration Routes of an Avian Herbivore [Barents Sea Data]

48. Descamps, S., et al. Data from: At-sea Distribution and Prey Selection of Antarctic Petrels and Commercial Krill Fisheries 2016.

49. Chudzińska, M. & Madsen, J. Data from: Foraging Behaviour and Fuel Accumulation of Capital Breeders during Spring Migration as Derived from a Combination of Satellite- and Ground-Based Observations 2016.

50. Herńandez-Pliego, J., Rodriguez, C. & Bustamante, J. Data from: Why Do Kestrels Soar? 2015.

51. Kölzsch, A., Kruckenberg, H., Glazov, P., Müskens, G. & Wikelski, M. Data from: Towards a New Understanding of Migration Timing: Slower Spring than Autumn Migration in Geese Reflects Different Decision Rules for Stopover Use and Departure 2016.

52. Arlt, D., Olsson, P., Fox, J. W., Low, M. & Pärt, T. Prolonged Stopover Duration Characterises Migration Strategy and Constraints of a Long-Distance Migrant Songbird. Animal Migration 2, 47–62. issn: 2084-8838 (Jan. 2015).

53. Bengtsson, D. et al. Does Influenza A Virus Infection Affect Movement Behaviour during Stopover in Its Wild Reservoir Host? Royal Society Open Science 3, 150633.

54. Bengtsson, D. et al. Movements, Home-Range Size and Habitat Selection of Mallards during Autumn Migration. PLOS ONE 9, e100764. issn: 1932-6203 (June 2014).

55. Boyd, W. S., Ward, D. H., Kraege, D. K. & Gerick, A. A. Migration Patterns of Western High Arctic (Grey-belly) Brant Branta Bernicla. Wildfowl 3, 3–25 (2013).

56. Bravo, S. P., Cueto, V. R. & Gorosito, C. A. Migratory Timing, Rate, Routes and Wintering Areas of White-crested Elaenia (Elaenia Albiceps Chilensis), a Key Seed Disperser for Patagonian Forest Regeneration. PLOS ONE 12, e0170188. issn: 1932-6203 (Feb. 2017).

57. Chudzińska, M. E., Nabe-Nielsen, J., Nolet, B. A. & Madsen, J. Foraging Behaviour and Fuel Accumulation of Capital Breeders during Spring Migration as Derived from a Combination of Satellite- and Ground-Based Observations. Journal of Avian Biology 47, 563–574. issn: 1600-048X (2016).

58. Cochran, W. & Wikelski, M. in Birds of Two Worlds: The Ecology and Evolution of Migration (The Johns Hopkins University Press, Baltimore, 2005). isbn: 978-0-8018-8107-7.

59. Cochran, W. W., Bowlin, M. S. & Wikelski, M. Wingbeat Frequency and Flap-Pause Ratio during Natural Migratory Flight in Thrushes. Integrative and Comparative Biology 48, 134–151. issn: 1540-7063 (July 2008).

60. DeLuca, W. V. et al. Transoceanic Migration by a 12 g Songbird. Biology Letters 11, 20141045 (Apr. 2015).

61. Descamps, S. et al. At-Sea Distribution and Prey Selection of Antarctic Petrels and Commercial Krill Fisheries. PLOS ONE 11, e0156968. issn: 1932-6203 (Aug. 2016).

62. van Toor, M., Ottosson, U., van der Meer, T., van Hoorn, S. & Waldenström, J. Data from: As the duck flies: estimating the dispersal of low-pathogenic avian influenza viruses by migrating mallards 2018. http://dx.doi.org/10.5441/001/1.3fv21n7m.

63. Efrat, R., Harel, R., Alexandrou, O., Catsadorakis, G. & Nathan, R. Seasonal Differences in Energy Expenditure, Flight Characteristics and Spatial Utilization of Dalmatian Pelicans Pelecanus Crispus in Greece. Ibis 161, 415–427. issn: 1474-919X (2019).

64. Efrat, R., Harel, R., Alexandrou, O., Catsadorakis, G. & Nathan, R. Seasonal Differences in Energy Expenditure, Flight Characteristics and Spatial Utilization of Dalmatian Pelicans *Pelecanus Crispus* in Greece. Ibis 161, 415–427. issn: 0019-1019, 1474-919X (Apr. 2019).

65. Eichhorn, G. in Seeking Nature’s Limits: Ecologists in the Field trans. by Drent, R. H., 84–90 (KNNV Pub., Utrecht, 2005). isbn: 978-90-5011-221-5.

66. Eichhorn, G. Travels in a Changing World Flexibility and Constraints in Migration and Breeding of the Barnacle Goose Thesis Fully Internal (DIV) (s.n., 2008). isbn: 9789036734486.

67. Flack, A. et al. Costs of Migratory Decisions: A Comparison across Eight White Stork Populations. Science Advances 2, e1500931 (Jan. 2016).

68. Flack, A., Nagy, M., Fiedler, W., Couzin, I. D. & Wikelski, M. From Local Collective Behavior to Global Migratory Patterns in White Storks. Science 360, 911–914 (May 2018).

69. Fleishman, E. et al. Space Use by Swainson’s Hawk (Buteo Swainsoni) in the Natomas Basin, California. Collabra 2, 5. issn: 2376-6832 (Apr. 2016).

70. Friedemann, G. et al. Multidimensional Differentiation in Foraging Resource Use during Breeding of Two Sympatric Top Predators. Scientific Reports 6, 35031. issn: 2045-2322 (Oct. 2016).

71. Fuller, M. R., Seegar, W. S. & Schueck, L. S. Routes and Travel Rates of Migrating Peregrine Falcons Falco Peregrinus and Swainson’s Hawks Buteo Swainsoni in the Western Hemisphere. Journal of Avian Biology 29, 433–440. issn: 0908-8857 (1998).

72. Hallworth, M. T. & Marra, P. P. Miniaturized GPS Tags Identify Non-breeding Territories of a Small Breeding Migratory Songbird. Scientific Reports 5, 11069. issn: 2045-2322 (June 2015).

73. Cueto, V. & Bravo, S. Data from: Migratory Timing, Rate, Routes and Wintering Areas of White-Crested Elaenia (Elaenia Albiceps Chilensis), a Key Seed Disperser for Patagonian Forest Regeneration 2017.

74. Harrison, A.-L., Woodard, P., Mallory, M. & Rausch, J. *Sympatrically-Breeding Congeneric Seabirds (Stercorarius Spp.) from Arctic Canada Migrate to Four Oceans* 2021.

75. Harrison, A.-L., Woodard, P. F., Mallory, M. L. & Rausch, J. Sympatrically Breeding Congeneric Seabirds (Stercorarius Spp.) from Arctic Canada Migrate to Four Oceans. Ecology and Evolution 12, e8451. issn: 2045-7758 (2022).

76. Herńandez-Pliego, J., Rodríguez, C. & Bustamante, J. Why Do Kestrels Soar? PLOS ONE 10, e0145402. issn: 1932-6203 (Dec. 2015).

77. Klein, K. et al. Fly with the Flock: Immersive Solutions for Animal Movement Visualization and Analytics. Journal of The Royal Society Interface 16, 20180794 (Apr. 2019).

78. Kleyheeg, E. et al. Movement Patterns of a Keystone Waterbird Species Are Highly Predictable from Landscape Configuration. Movement Ecology 5, 2. issn: 2051-3933 (Feb. 2017).

79. Kochert, M. N. et al. Migration Patterns, Use of Stopover Areas, and Austral Summer Movements of Swainson’s Hawks. The Condor 113, 89–106. issn: 1938-5129 (Feb. 2011).

80. Kölzsch, A., et al. Forecasting Spring from Afar? Timing of Migration and Predictability of Phenology along Different Migration Routes of an Avian Herbivore. Journal of Animal Ecology 84, 272–283. issn: 1365-2656 (2015).

81. Kölzsch, A., et al. Towards a New Understanding of Migration Timing: Slower Spring than Autumn Migration in Geese Reflects Different Decision Rules for Stopover Use and Departure. Oikos 125, 1496–1507. issn: 1600-0706 (2016).

82. Köppen, U., Yakovlev, A. P., Barth, R., Kaatz, M. & Berthold, P. Seasonal Migrations of Four Individual Bar-Headed Geese Anser Indicus from Kyrgyzstan Followed by Satellite Telemetry. Journal of Ornithology 151, 703–712. issn: 2193-7206 (July 2010).

83. Korner, P., Sauter, A., Fiedler, W. & Jenni, L. Variable Allocation of Activity to Daylight and Night in the Mallard. Animal Behaviour 115, 69–79. issn: 0003-3472 (May 2016).

84. Ross, J., Bridge, E., Rozmarynowycz, M. & Bingman, V. Data from: Individual Variation in Migratory Path and Behavior among Eastern Lark Sparrows 2014.

85. Lamb, J. S., Satgé, Y. G. & Jodice, P. G. R. Influence of Density-Dependent Competition on Foraging and Migratory Behavior of a Subtropical Colonial Seabird. Ecology and Evolution 7, 6469–6481. issn: 2045-7758 (2017).

86. Lyons, D. E., Patterson, A. G. L., Tennyson, J., Lawes, T. J. & Roby, D. D. The Salton Sea: Critical Migratory Stopover Habitat for Caspian Terns (Hydroprogne Caspia) in the North American Pacific Flyway. Waterbirds 41, 154–165. issn: 1524-4695, 1938-5390 (June 2018).

87. Nagy, M., Couzin, I. D., Fiedler, W., Wikelski, M. & Flack, A. Synchronization, Coordination and Collective Sensing during Thermalling Flight of Freely Migrating White Storks. Philosophical Transactions of the Royal Society B: Biological Sciences 373, 20170011 (May 2018).

88. Petersen, M. R., Earned, W. W. & Douglas, D. C. At-Sea Distribution of Spectacled Eiders: A 120-Year-Old Mystery Resolved. The Auk 116, 1009–1020. issn: 1938-4254 (Oct. 1999).

89. Petersen, M. R., Grand, J. B. & Dau, C. P. Spectacled Eider (Somateria Fischeri), Version 2.0. Birds of North America (2000).

90. Petersen, M., Flint, P. L., Mulcahy, D. & Douglas, D. C. Tracking Data for Common Eiders (Somateria Mollissima) 2021.

91. Petersen, M. R., Douglas, D. C. & Mulcahy, D. M. Use of Implanted Satellite Transmitters to Locate Spectacled Eiders At-Sea. The Condor 97, 276–278. issn: 0010-5422 (1995).

92. Petersen, M. R. & Douglas, D. C. Winter Ecology of Spectacled Eiders: Environmental Characteristics and Population Change. The Condor 106, 79–94. issn: 0010-5422 (2004).

93. Poli, C. L., Harrison, A.-L., Vallarino, A., Gerard, P. D. & Jodice, P. G. R. Dynamic Oceanography Determines Fine Scale Foraging Behavior of Masked Boobies in the Gulf of Mexico. PLOS ONE 12, e0178318. issn: 1932-6203 (June 2017).

94. Ross, J. D., Bridge, E. S., Rozmarynowycz, M. J. & Bingman, V. P. Individual Variation in Migratory Path and Behavior among Eastern Lark Sparrows. Animal Migration 2, 29–33. issn: 2084-8838 (Jan. 2015).

95. Fleishman, E., et al. Data from: Space Use by Swainson’s Hawk (Buteo Swainsoni) in the Natomas Basin, California

96. Senner, N. R. et al. When Siberia Came to the Netherlands: The Response of Continental Black-Tailed Godwits to a Rare Spring Weather Event. Journal of Animal Ecology 84, 1164–1176. issn: 1365-2656 (2015).

97. Sexson, M. G., Petersen, M. R., Breed, G. A. & Powell, A. N. Shifts in the Distribution of Molting Spectacled Eiders (Somateria Fischeri) Indicate Ecosystem Change in the Arctic. The Condor 118, 463–476. issn: 1938-5129 (Aug. 2016).

98. Shariati-Najafabadi, M. et al. Environmental Parameters Linked to the Last Migratory Stage of Barnacle Geese En Route to Their Breeding Sites. Animal Behaviour 118, 81–95. issn: 0003-3472 (Aug. 2016).

99. Shariatinajafabadi, M. et al. Migratory Herbivorous Waterfowl Track Satellite-Derived Green Wave Index. PLOS ONE 9, e108331. issn: 1932-6203 (Sept. 2014).

100. Shariati Najafabadi, M., et al. Satellite-versus Temperature-Derived Green Wave Indices for Predicting the Timing of Spring Migration of Avian Herbivores. Ecological Indicators 58, 322–331. issn: 1470-160X (Nov. 2015).

101. Silveira, N. S. D., Niebuhr, B. B. S., Muylaert, R. d. L., Ribeiro, M. C. & Pizo, M. A. Effects of Land Cover on the Movement of Frugivorous Birds in a Heterogeneous Landscape. PLOS ONE 11, e0156688. issn: 1932-6203 (June 2016).

102. Soanes, L. M., Atkinson, P. W., Gauvain, R. D. & Green, J. A. Individual Consistency in the Foraging Behaviour of Northern Gannets: Implications for Interactions with Offshore Renewable Energy Developments. Marine Policy 38, 507–514. issn: 0308-597X (Mar. 2013).

103. Spiegel, O., Getz, W. M. & Nathan, R. Factors Influencing Foraging Search Efficiency: Why Do Scarce Lappet-Faced Vultures Outperform Ubiquitous White-Backed Vultures? The American Naturalist 181, E102–E115. issn: 0003-0147 (May 2013).

104. Tarroux, A. et al. Flexible Flight Response to Challenging Wind Conditions in a Commuting Antarctic Seabird: Do You Catch the Drift? Animal Behaviour 113, 99–112. issn: 0003-3472 (Mar. 2016).

105. Van Toor, M. L. et al. Flexibility of Continental Navigation and Migration in European Mallards. PLOS ONE 8, e72629. issn: 1932-6203 (Aug. 2013).

106. Flack, A., et al. Data from: Costs of Migratory Decisions: A Comparison across Eight White Stork Populations 2015.

107. Torres, L. G., Orben, R. A., Tolkova, I. & Thompson, D. R. Classification of Animal Movement Behavior through Residence in Space and Time. PLOS ONE 12, e0168513. issn: 1932-6203 (Jan. 2017).

108. Tucker, M. A. et al. Large Birds Travel Farther in Homogeneous Environments. Global Ecology and Biogeography 28 (ed Boucher-Lalonde, V.) 576–587. issn: 1466-822X, 1466-8238 (May 2019).

109. van Toor, M. L., Avril, A., Wu, G., Holan, S. H. & Waldenström, J. As the Duck Flies—Estimating the Dispersal of Low-Pathogenic Avian Influenza Viruses by Migrating Mallards. Frontiers in Ecology and Evolution 6. issn: 2296-701X (2018).

110. Warwick-Evans, V. et al. Changes in Behaviour Drive Inter-Annual Variability in the at-Sea Distribution of Northern Gannets. Marine Biology 163, 156. issn: 1432-1793 (June 2016).

111. Warwick-Evans, V., Atkinson, P. W., Walkington, I. & Green, J. A. Predicting the Impacts of Wind Farms on Seabirds: An Individual-Based Model. Journal of Applied Ecology 55, 503–515. issn: 1365-2664 (2018).

112. Weinzierl, R. et al. Wind Estimation Based on Thermal Soaring of Birds. Ecology and Evolution 6, 8706–8718. issn: 2045-7758 (2016).

113. Wikelski, M. et al. True Navigation in Migrating Gulls Requires Intact Olfactory Nerves. Scientific Reports 5, 17061. issn: 2045-2322 (Nov. 2015).

114. Ens, B. et al. *SOVON-onderzoeksrapport 2008/*10: Tracking of individual birds. Report on WP 3230 (bird tracking sensor characterization) and WP 4130 (sensor adaptation and calibration for bird tracking system) of the FlySafe basic activities project. English. Reporting year: 2008 (SOVON Vogelonderzoek Nederland, Netherlands, 2008).

115. DeLuca, W., et al. Data from: Transoceanic migration by a 12 g songbird 2015. http://dx.doi.org/10.5441/001/1.jb182ng4.

116. Georgopoulou, E., Alexandrou, O., Manolopoulos, A., Xirouchakis, S. & Catsadorakis, G. Home Range of the Dalmatian Pelican in South-East Europe. European Journal of Wildlife Research 69, 41. issn: 1439-0574. https://doi.org/10.1007/s10344-023-01667-1 (2023) (Mar. 31, 2023).

117. Lamb, J., Satgé, Y. & Jodice, P. Data from: Influence of Density-Dependent Competition on Foraging and Migratory Behavior of a Subtropical Colonial Seabird 2017.

118. Scott, T. Movements of White-Headed and White-Backed Vultures. Boise State University Theses and Dissertations. https://scholarworks.boisestate.edu/td/1716 (Aug. 1, 2020).

119. Matthes, D., et al. Data from: Flexibility of Continental Navigation and Migration in European Mallards 2013.

120. Zhang, G. et al. Bidirectional Movement of Emerging H5N8 Avian Influenza Viruses Between Europe and Asia via Migratory Birds Since Early 2020. Molecular Biology and Evolution 40, msad019. issn: 1537-1719. https://doi.org/10.1093/molbev/msad019 (2023) (Feb. 1, 2023).

121. Chen, T. & Guestrin, C. XGBoost: A Scalable Tree Boosting System in Proceedings of the 22nd ACM SIGKDD International Conference on Knowledge Discovery and Data Mining (Association for Computing Machinery, New York, NY, USA, Aug. 2016), 785–794. isbn: 978-1-4503-4232-2. https://doi.org/10.1145/2939672.2939785 (2020).

122. Van Doren, B. M. & Horton, K. G. A continental system for forecasting bird migration. Science 361, 1115–1118. issn: 0036-8075. http://www.sciencemag.org/lookup/doi/10.1126/science.aat7526 (Sept. 2018).

123. Shu, Y. & McCauley, J. GISAID: Global initiative on sharing all influenza data – from vision to reality. en. Eurosurveillance 22. Publisher: European Centre for Disease Prevention and Control, 30494. issn: 1560-7917. https://www.eurosurveillance.org/content/10.2807/1560-7917.ES.2017.22.13.30494?crawler=true (2022) (Mar. 2017).

124. Khare, S. et al. GISAID’s Role in Pandemic Response. China CDC Weekly 3, 1049–1051. issn: 2096-7071 (Dec. 2021).

125. Elbe, S. & Buckland-Merrett, G. Data, Disease and Diplomacy: GISAID’s Innovative Contribution to Global Health. Global Challenges 1, 33–46. issn: 2056-6646 (2017).

126. Price, M., Dehal, P. & Arkin, A. FastTree: computing large minimum evolution trees with profiles instead of a distance matrix. en. Mol Biol Evol 26, 1641–1650 (2009).

127. Lam, T.-Y. The genesis and source of the H7N9 influenza viruses causing human infections in China. en. Nature 502, 241 (2013).

128. Rambaut, A., Lam, T., Carvalho, L. & Pybus, O. Exploring the temporal structure of heterochronous sequences using TempEst (formerly Path-O-Gen. en. Virus Evol 2, vew007 (2016).

129. Suchard, M. Bayesian phylogenetic and phylodynamic data integration using BEAST 1.10. en. Virus Evolution 4 (2018).

130. Ayres, D. BEAGLE: an application programming interface and high-performance computing library for statistical phylogenetics. en. Syst Biol syr100 (2011).

131. Drummond, A., Ho, S., Phillips, M. & Rambaut, A. Relaxed phylogenetics and dating with confidence. en. PLoS Biol 4, 88 (2006).

132. Shapiro, B., Rambaut, A. & Drummond, A. Choosing Appropriate Substitution Models for the Phylogenetic Analysis of Protein-Coding Sequences. en. Mol Biol Evol 23, 7–9 (2005).

133. Minin, V., Bloomquist, E. & Suchard, M. Smooth skyride through a rough skyline: Bayesian coalescent-based inference of population dynamics. en. Mol Biol Evol 25, 1459–1471 (2008).

134. Rambaut, A., Drummond, A., Xie, D., Baele, G. & Suchard, M. Posterior Summarization in Bayesian Phylogenetics Using Tracer 1.7. cs. Systematic Biology 67, 901–904 (2018).

135. O’Brien, J. D., Minin, V. N. & Suchard, M. A. Learning to Count: Robust Estimates for Labeled Distances between Molecular Sequences. Molecular Biology and Evolution 26, 801–814. issn: 0737-4038. https://doi.org/10.1093/molbev/msp003 (2021) (Apr. 2009).

136. Minin, V. N. & Suchard, M. A. Fast, accurate and simulation-free stochastic mapping. Philosophical Transactions of the Royal Society B: Biological Sciences 363. Publisher: Royal Society, 3985–3995. https://royalsocietypublishing.org/doi/10.1098/rstb.2008.0176 (2021) (Dec. 2008).

137. Faria, N. R., Suchard, M. A., Rambaut, A., Streicker, D. G. & Lemey, P. Simultaneously reconstructing viral crossspecies transmission history and identifying the underlying constraints. Philosophical Transactions of the Royal Society B: Biological Sciences 368. issn: 14712970 (2013).

138. Faria, N. R., et al. Distinct rates and patterns of spread of the major HIV-1 sub-types in Central and East Africa. PLoS Pathogens 15. Publisher: Public Library of Science, e1007976. issn: 15537374 (2019).

139. Faria, N. R. et al. The early spread and epidemic ignition of HIV-1 in human populations. Science 346, 56–61. issn: 10959203 (2014).

140. Vasylyeva, T. I. et al. Molecular epidemiology reveals the role of war in the spread of HIV in Ukraine. Proceedings of the National Academy of Sciences of the United States of America 115, 1051–1056. issn: 10916490 (2018).

141. Thézé, J., et al. Genomic Epidemiology Reconstructs the Introduction and Spread of Zika Virus in Central America and Mexico. Cell Host and Microbe 23, 855– 864.e7. issn: 19346069 (2018).

142. Yang, Q. et al. Assessing the role of live poultry trade in community-structured transmission of avian influenza in China. en. Proceedings of the National Academy of Sciences 117, 5949–5954. issn: 0027-8424, 1091-6490. http://www.pnas.org/lookup/doi/10.1073/pnas.1906954117 (2021) (Mar. 2020).

