## Supplemental materials for "Synchrony of Bird Migration with Avian Influenza Global Spread; Implications for Vulnerable Bird Orders"

**This PDF file includes:**

Figure. S1 to S7
Table S2, S4, S5
References

**Other Supplementary Material for this manuscript includes the following:**

Table S1, 3, 6 as Excel files
Analysis scripts (BEAST xml files, Python/R files): GitHub repository
https://github.com/kikiyang/HPAI_Bird_world

**Table S2** Independent variables of the species distribution model of wild birds.

| Variables | Source |
|---|---|
| Normalized Difference Vegetation Index monthly mean | Pinzon et al(2014) [25] |
| Evergreen Deciduous Needleleaf Trees | Tuanmu and Jetz (2014) [26] |
| Evergreen Broadleaf Trees | Tuanmu and Jetz (2014) [26] |
| Deciduous Broadleaf Trees | Tuanmu and Jetz (2014) [26] |
| Mixed/Other Trees | Tuanmu and Jetz (2014) [26] |
| Shrubs | Tuanmu and Jetz (2014) [26] |
| Herbaceous Vegetation | Tuanmu and Jetz (2014) [26] |
| Cultivated and Managed Vegetation | Tuanmu and Jetz (2014) [26] |
| Regularly Flooded Vegetation | Tuanmu and Jetz (2014) [26] |
| Urban/Built-up | Tuanmu and Jetz (2014) [26] |
| Snow-Ice | Tuanmu and Jetz (2014) [26] |
| Barren | Tuanmu and Jetz (2014) [26] |
| Open Water | Tuanmu and Jetz (2014) [26] |
| Mean Temperature | WorldClim 2.1[27] |
| Wind Speed | WorldClim 2.1[27] |
| Total Precipitation | WorldClim 2.1[27] |
| Elevation | SRTM elevation data[28–32] |

**Table S4.1** Bird orders at the origin location whose distribution probability significantly correlates (*P* < 0.00556) with virus lineage movements from the origin to the destination location, leading by some month lags. The confidence interval of the correlation is (-0.566, 0.566).

| **Bird order** | **Origin** | **Destination** | **Month lags** | **Correlation** | ***P*-value** |
|---|---|---|---|---|---|
| Charadriiformes | SChina | NChina | -1 | 0.816 | 0.005 |
| Charadriiformes | Qinghai | Europe | 0 | -0.820 | 0.004 |
| Ciconiiformes | NChina | Europe | -3 | 0.845 | 0.003 |
| Suliformes | JapanKorea | Europe | -2 | 0.863 | 0.003 |
| Falconiformes | JapanKorea | Europe | -2 | 0.804 | 0.005 |
| Pelecaniformes | NChina | JapanKorea | 0 | 0.818 | 0.005 |
| Ciconiiformes | NChina | JapanKorea | 0 | 0.952 | 0.001 |
| Ciconiiformes | NChina | Qinghai | -3 | 0.808 | 0.005 |
| Charadriiformes | SChina | SEA | 0 | 0.803 | 0.005 |
| Suliformes | Europe | Qinghai | 0 | 0.966 | 0.001 |
| Ciconiiformes | Europe | Africa | -3 | 0.813 | 0.005 |

**Table S4.2** Bird orders at the destination location whose distribution probability has significantly positive correlation (*P* < 0.00556) with virus lineage movements from the origin to the destination location, lagging by some month lags. The confidence interval of the correlation is (-0.566, 0.566).

| **Bird order** | **Origin** | **Destination** | **Month lags** | **Correlation** | ***P*-value** |
|---|---|---|---|---|---|
| Passeriformes | Qinghai | Europe | 1 | 0.878 | 0.002 |
| Suliformes | Qinghai | Europe | 0 | 0.924 | 0.001 |
| Accipitriformes | JapanKorea | USAandCanada | 0 | 0.922 | 0.001 |
| Suliformes | SChina | SEA | 0 | 0.912 | 0.002 |
| Suliformes | SChina | SEA | 1 | 0.820 | 0.004 |
| Falconiformes | SChina | SEA | 0 | 0.890 | 0.002 |
| Suliformes | Europe | Qinghai | 2 | 0.899 | 0.002 |
| Anseriformes | NChina | SChina | 0 | 0.827 | 0.004 |
| Charadriiformes | Europe | Africa | 1 | 0.886 | 0.002 |
| Anseriformes | Europe | Africa | 1 | 0.905 | 0.002 |
| Charadriiformes | Russia | Africa | 1 | 0.881 | 0.002 |
| Anseriformes | Russia | Africa | 1 | 0.937 | 0.001 |

**Table S5** Macaulay Library entries of the bird species photos in Figure 2.

| Species | Order | Entry |
|---|---|---|
| *Grus grus* | *Gruiformes* | ML113809411 |
| *Phalacrocoracidae sp.* | *Suliformes* | ML488694141 |
| *Sylvia atricapilla* | *Passeriformes* | ML466144251 |
| *Cygnus olor* | *Anseriformes* | ML72775261 |
| *Pelecanus crispus* | *Pelecaniformes* | ML461113431 |
| *Ciconia ciconia* | *Ciconiiformes* | ML41115011 |
| *Tringa totanus* | *Charadriiformes* | ML466721131 |
| *Pandion haliaetus haliaetus* | *Accipitriformes* | ML523577271 |
| *Falco amurensis* | *Falconiformes* | ML538883721 |

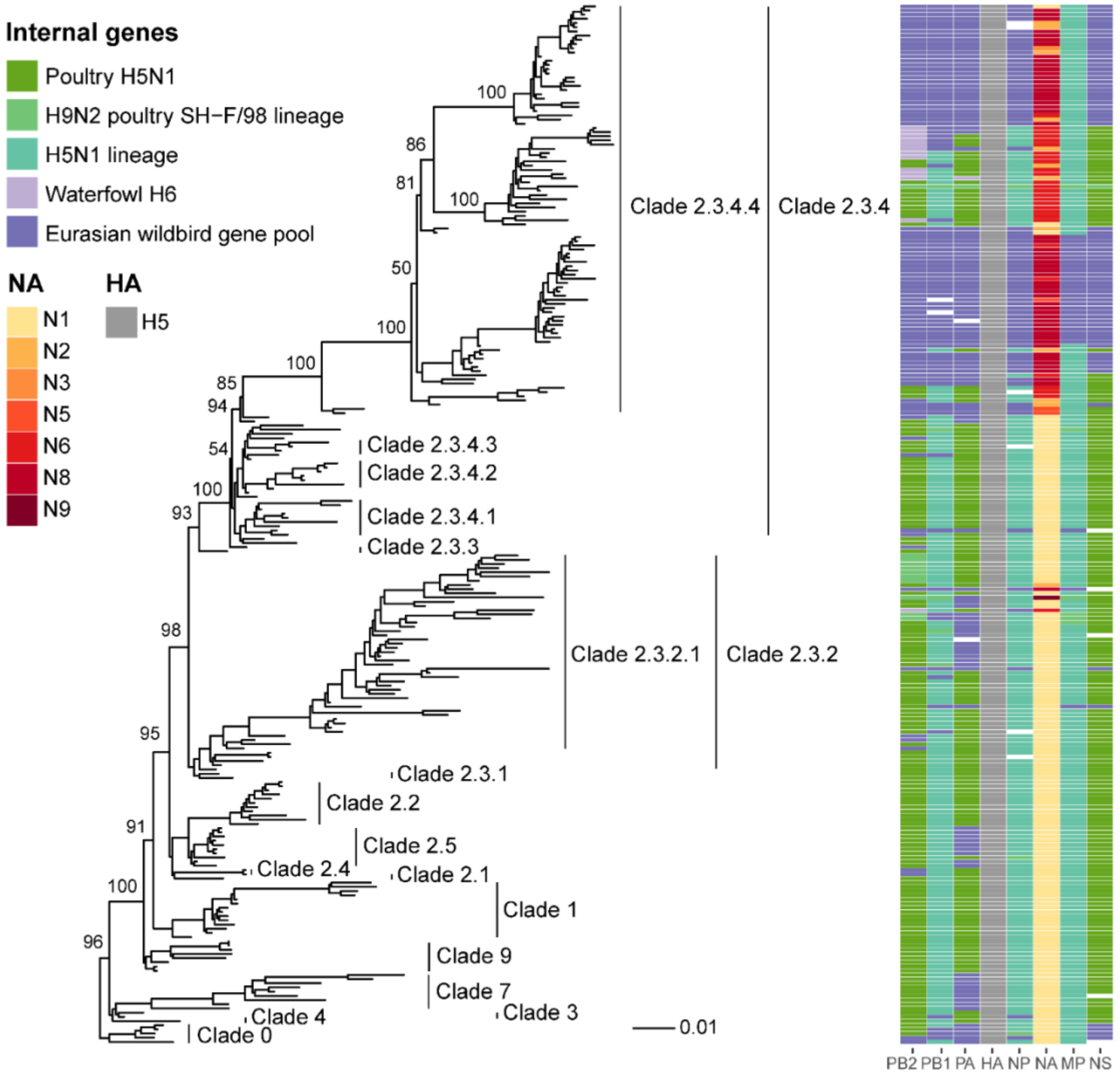


**Figure S3.** Maximum likelihood tree of hemagglutinin (HA) genes and genotypes of all gene segments of H5 subtype avian influenza A viruses.

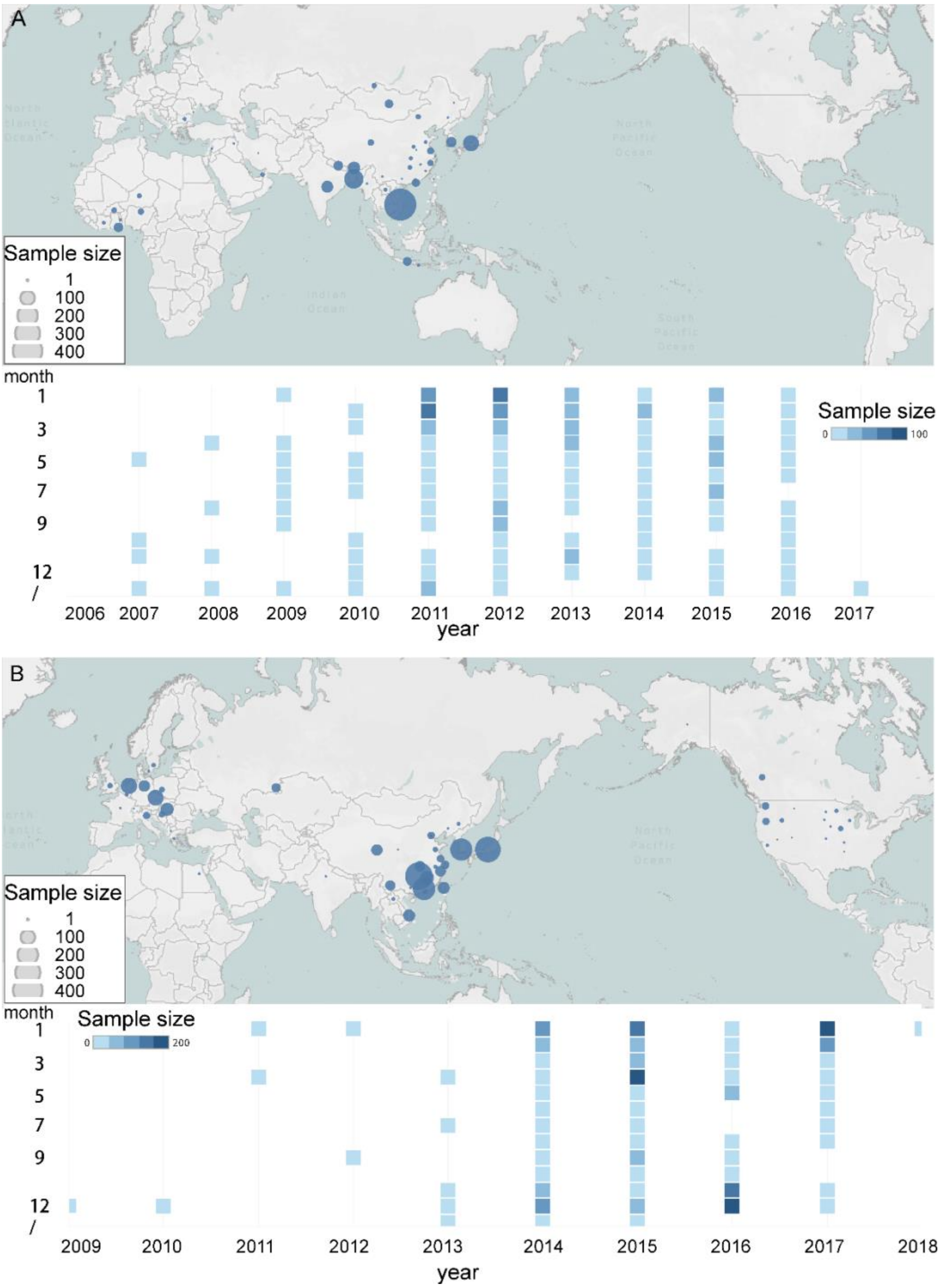


**Figure S4** Spatial-temporal distribution of HA sequences of Clade 2.3.2.1 ***(A)*** and Clade 2.3.4.4 ***(B)***.

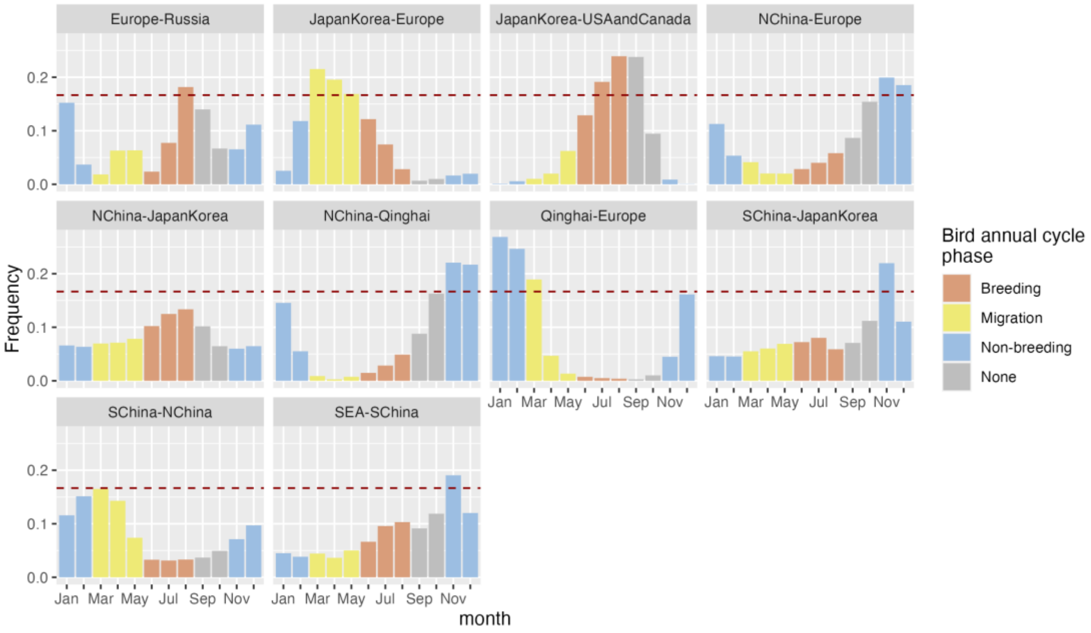


**Figure S5.1** Frequency density distribution of Markov jump events of clade 2.3.4.4 each month through a year from southern to northern regions. We consider a peak of virus lineage movement when the frequency > 0.167 (=2*0.083, if the frequency is evenly distribution across months, monthly frequency should be 0.083).

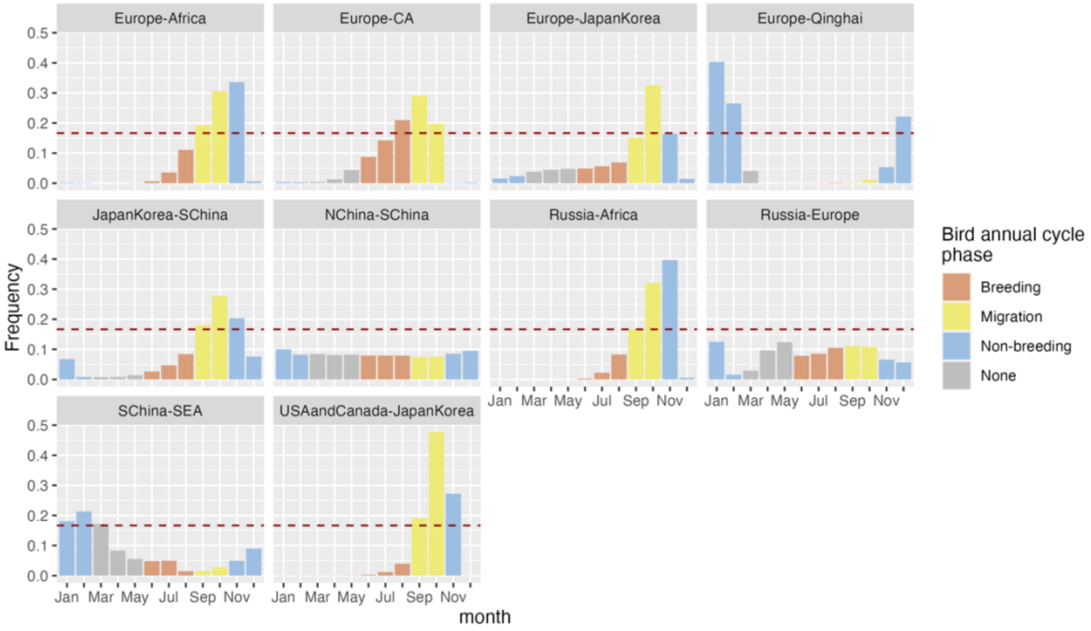


**Figure S5.2** Frequency density distribution of Markov jump events of clade 2.3.4.4 each month through a year from year from northern to southern regions. We consider a peak of virus lineage movement when the frequency > 0.167 (=2*0.083, if the frequency is evenly distribution across months, monthly frequency should be 0.083).

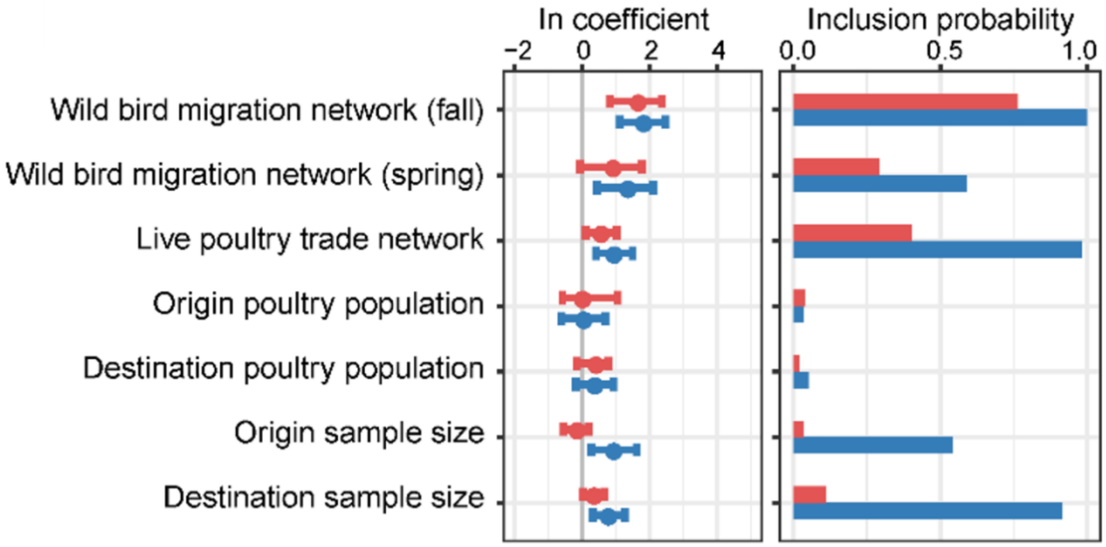


**Figure S6** Contributions of predictors to worldwide diffusion of Clade 2.3.2.1 and Clade 2.3.4.4 of H5N1 inferred on HA genes by GLM-extended Bayesian phylogeographic inference heterogeneous evolutionary processes through time. Spring and migration networks of wild birds are two separate indicators in the model.

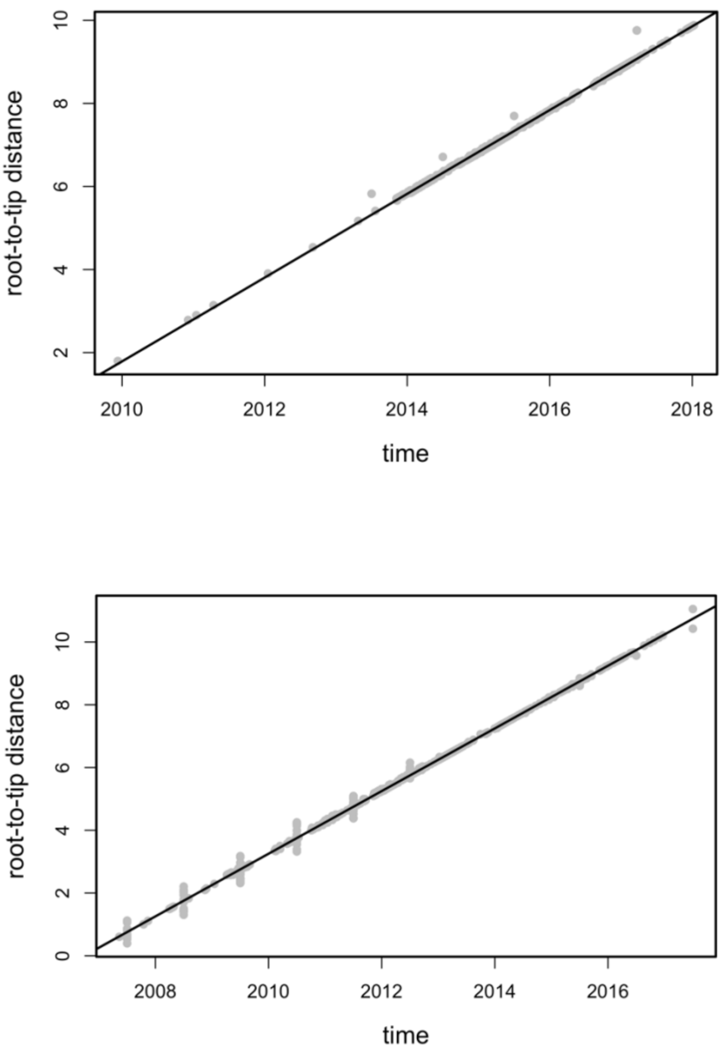


**Figure S7** Strong temporal signal tested in TempEst of HA genes of HPAIV H5 clade 2.3.4.4 (top panel) and clade 2.3.2.1 (bottom panel) (both R squared > 0.99). We used the final dataset of 1844 HA sequences of clade 2.3.4.4 and 1163 HA sequences of clade 2.3.2.1.

### References


1. Chen, T. & Guestrin, C. XGBoost: A Scalable Tree Boosting System. in *Proceedings of the 22nd ACM SIGKDD International Conference on Knowledge Discovery and Data Mining* 785–794 (Association for Computing Machinery, 2016). doi:10.1145/2939672.2939785.
2. Price, M. N., Dehal, P. S. & Arkin, A. P. FastTree: computing large minimum evolution trees with profiles instead of a distance matrix. *Mol Biol Evol* **26**, 1641–1650 (2009).
3. Lam, T. T. Y. *et al.* The genesis and source of the H7N9 influenza viruses causing human infections in China. *Nature* **502**, 241–244 (2013).
4. Rambaut, A., Lam, T. T., Max Carvalho, L. & Pybus, O. G. Exploring the temporal structure of heterochronous sequences using TempEst (formerly Path-O-Gen). *Virus Evolution* **2**, vew007 (2016).
5. Suchard, M. A. Bayesian phylogenetic and phylodynamic data integration using BEAST 1.10. *Virus Evolution* **4**, (2018).
6. Ayres, D. L. BEAGLE: an application programming interface and high-performance computing library for statistical phylogenetics. *Syst Biol* **syr100**, (2011).
7. Drummond, A. J., Ho, S. Y., Phillips, M. J. & Rambaut, A. Relaxed phylogenetics and dating with confidence. *PLoS Biol* **4**, 88 (2006).
8. Shapiro, B., Rambaut, A. & Drummond, A. J. Choosing Appropriate Substitution Models for the Phylogenetic Analysis of Protein-Coding Sequences. *Mol Biol Evol* **23**, 7–9 (2005).

9. Minin, V. N., Bloomquist, E. W. & Suchard, M. A. Smooth skyride through a rough skyline: Bayesian coalescent-based inference of population dynamics. *Mol Biol Evol* **25**, 1459–1471 (2008).
10. Rambaut, A., Drummond, A. J., Xie, D., Baele, G. & Suchard, M. A. Posterior Summarization in Bayesian Phylogenetics Using Tracer 1.7. *Systematic Biology* **67**, 901–904 (2018).
11. Lemey, P., Rambaut, A., Drummond, A. J. & Suchard, M. A. Bayesian Phylogeography Finds Its Roots. *PLoS Computational Biology* **5**, e1000520 (2009).
12. O'Brien, J. D., Minin, V. N. & Suchard, M. A. Learning to Count: Robust Estimates for Labeled Distances between Molecular Sequences. *Molecular Biology and Evolution* **26**, 801–814 (2009).
13. Minin, V. N. & Suchard, M. A. Fast, accurate and simulation-free stochastic mapping. *Philosophical Transactions of the Royal Society B: Biological Sciences* **363**, 3985–3995 (2008).
14. Faria, N. R., Suchard, M. A., Rambaut, A., Streicker, D. G. & Lemey, P. Simultaneously reconstructing viral crossspecies transmission history and identifying the underlying constraints. *Philosophical Transactions of the Royal Society B: Biological Sciences* **368**, (2013).
15. Faria, N. R. *et al.* Distinct rates and patterns of spread of the major HIV-1 subtypes in Central and East Africa. *PLoS Pathogens* **15**, e1007976 (2019).

16. Faria, N. R. *et al.* The early spread and epidemic ignition of HIV-1 in human populations. *Science* **346**, 56–61 (2014).
17. Vasylyeva, T. I. *et al.* Molecular epidemiology reveals the role of war in the spread of HIV in Ukraine. *Proceedings of the National Academy of Sciences of the United States of America* **115**, 1051–1056 (2018).
18. Thézé, J. *et al.* Genomic Epidemiology Reconstructs the Introduction and Spread of Zika Virus in Central America and Mexico. *Cell Host and Microbe* **23**, 855-864.e7 (2018).
19. Bielejec, F. SpreaD3: Interactive Visualization of Spatiotemporal History and Trait Evolutionary Processes. *Molecular biology and evolution* **33**, 2167–2169 (2016).
20. Lemey, P. *et al.* Unifying Viral Genetics and Human Transportation Data to Predict the Global Transmission Dynamics of Human Influenza H3N2. *PLoS pathogens* **10**, 1–17 (2014).
21. Bielejec, F., Lemey, P., Baele, G., Rambaut, A. & Suchard, M. A. Inferring heterogeneous evolutionary processes through time: From sequence substitution to phylogeography. *Systematic Biology* **63**, 493–504 (2014).
22. Yang, Q. *et al.* Assessing the role of live poultry trade in community-structured transmission of avian influenza in China. *Proceedings of the National Academy of Sciences of the United States of America* **117**, 5949–5954 (2020).
23. Burt, W. H. Territoriality and Home Range Concepts as Applied to Mammals. *Journal of Mammalogy* **24**, 346–352 (1943).

24. Kernel Methods for Estimating the Utilization Distribution in Home-Range Studies on JSTOR. https://www.jstor.org/stable/1938423?seq=1#metadata_info_tab_contents.

25. Pinzon, J. E. & Tucker, C. J. A Non-Stationary 1981–2012 AVHRR NDVI3g Time Series. *Remote Sensing* **6**, 6929–6960 (2014).

26. Tuanmu, M.-N. & Jetz, W. A global 1-km consensus land-cover product for biodiversity and ecosystem modelling. *Global Ecology and Biogeography* **23**, 1031–1045 (2014).

27. Fick, S. E. & Hijmans, R. J. WorldClim 2: new 1-km spatial resolution climate surfaces for global land areas. *International Journal of Climatology* **37**, 4302–4315 (2017).

28. Farr, T. G. & Kobrick, M. Shuttle radar topography mission produces a wealth of data. *Eos, Transactions American Geophysical Union* **81**, 583–585 (2000).

29. Farr, T. G. *et al.* The Shuttle Radar Topography Mission. *Rev. Geophys.* **45**, RG2004 (2007).

30. OpenTopography. Shuttle Radar Topography Mission (SRTM) Global. (2013) doi:10.5069/G9445JDF.

31. Rosen, P. A. *et al.* Synthetic aperture radar interferometry. *Proceedings of the IEEE* **88**, 333–382 (2000).

32. Kobrick, M. On the Toes of Giants - How SRTM was Born. *Photogramm. Eng. Remote Sens.* 206–210 (2006).
